# Innovative platelet-derived extracellular vesicle drug delivery system for the treatment of corneal neovascularization

**DOI:** 10.1101/2024.09.25.614855

**Authors:** Guei-Sheung Liu, Huai-An Chen, Che-Yi Chang, Yin-Ju Chen, Yu-Yi Wu, Ariel Widhibrata, Ya-Han Yang, Erh-Hsuan Hsieh, Liling Delila, I-Chan Lin, Thierry Burnouf, Ching-Li Tseng

## Abstract

Platelet-derived extracellular vesicles (PEVs) have drawn attention due to their multifunctionality, ease of procurement, and abundant supply from clinical-grade platelet concentrates. PEVs can be easily endocytosed owing to their lipid bilayer membrane and nanosized structure, thereby increasing the bioavailability and functionality of their therapeutic effects. PEVs also possess multiple trophic factors that make them effective therapeutic agents. Since nanomedicine offers advantages over traditional therapies for eye diseases by overcoming physical ocular barriers, PEVs combined with an anti-angiogenic agent, kaempferol (KM), were evaluated for their ability to inhibit abnormal vessel formation in the cornea. Characterization of the nanoparticles indicated successful preparation of KM- loaded PEVs (PEV-KM) with a mean diameter of ∼160 nm and an encapsulation efficiency of ∼61%. PEV-KM was efficiently internalized into human vascular endothelial cells, resulting in inhibited function, evidenced by lower wound closure rates, reduced tube formation capacity, and downregulation of angiogenesis-related gene expression. Moreover, prolonged ocular retention was observed followed by topical application of PEV and PEV-KM in mouse eyes. In an alkali-burned corneal neovascularization (CoNV) mouse model, PEV (1%) was found to reduce vessel formation in the injured cornea. PEV with KM (1% PEV with KM 6 µg/mL) showed an even stronger effect in suppressing CoNV and reducing the expression of proangiogenic and inflammatory cytokines. Together, our data suggests that topical administration of PEVs or in combination with KM (PEV-KM) is a promising therapeutic for managing CoNV.

## 1. Introduction

Corneal neovascularization (CoNV) is a pathological condition induced by infection, inflammation, trauma, corneal degeneration, or hypoxia [1, 2]. The changes in the cornea – the growth of new vessels into the normally clear tissue – significantly affect visual acuity and quality, including threatening corneal transparency, reducing vision, and even causing blindness [3, 4]. Current treatment strategies for managing CoNV include corneal transplantation, photodynamic therapy, laser photocoagulation, fine-needle diathermy, subconjunctival injection, and topical steroidal eye drops [1, 3]. However, steroid-containing eye drops pose risks of elevating intraocular pressure and can cause cataracts and glaucoma after long-term use [5, 6]. Therapeutic agents against the vascular endothelial growth factor (VEGF), such as aflibercept and bevacizumab, have been clinically used in managing neovascular eye conditions, neovascular age-related macular degeneration and diabetic retinopathy which all involve over-expression of VEGF in their pathologies [7, 8]. These anti-VEGF agents have also been approved for treating CoNV [8, 9]. While it is effective, the clinical application of anti- VEGF agents in corneal diseases is also limited by costs, short half-lives, reduced epithelial recovery rate, and corneal thinning [10]. Therefore, an alternative therapeutic strategy for treating CoNV is demanded.

Nanoparticles (NPs) have been extensively explored as delivery vehicles for ocular conditions, including CoNV [11, 12]. Due to the size effect and surface properties of NPs, they can greatly improve drug retention and bioavailability in the eyes, enabling effective treatment through topical delivery. Extracellular vehicles (EVs) are also known as natural nanoparticles and are lipid bilayer membrane particles ranging in size from approximately 30 to 1000 nm. Some examples of natural NPs include exosomes, microvesicles, and apoptotic bodies, which are released by cells into the extracellular environment and serve as important mediators in cell-cell communication [13, 14]. EVs carry a variety of biomolecules, including proteins, lipids, and nucleic acids (DNA, mRNA, miRNA, and other non- coding RNAs), which reflect their cell of origin and play significant roles in physiological and pathological functions [15]. EVs have recently garnered considerable translational interest in nanomedicine [16, 17] due to their diverse cargo and the presence of various surface proteins that can mediate targeting. For example, membrane proteins, such as integrins and cell-specific proteins, can act as native targeting moieties [18, 19]. Furthermore, their natural ability to cross tissue barriers, infiltrate deeper tissues [18, 20], and evade endosomal or lysosomal degradation [21] may enhance drug delivery efficiency.

There is growing interest in evaluating blood cell-derived EVs, particularly platelet-derived EVs (PEV) as biotherapies of degenerative diseases and as drug delivery systems [20, 22]. PEVs are the most abundant (about 50% or more) of all EVs present in the blood circulation [20, 23]. Advantageously, PEVs can readily be isolated from clinical-grade human platelet concentrates (PC), which are licensed therapeutic cellular products on the list of essential medicines of the World Health Organization. PEVs express numerous membrane receptors (e.g. CD41, CD61, and CD62P) originating from the platelets, helping to target sites of inflammation and damaged vascular tissue [20, 24]. Additionally, PEVs are a potent functional reservoir of trophic factors, including fibroblast growth factor-2 (FGF-2), epidermal growth factor (EGF), platelet-derived growth factors (PDGF), transforming growth factor-beta (TGF-β), insulin-like growth factor-1 (IGF-1), VEGF and many more [25–28]. Growth factors such as EGF and IGF-1 improve wound healing by stimulating the migration, proliferation, and differentiation of keratinocytes or fibroblasts [29, 30]. Additionally, FGF2, PDGF, and TGF-β contained in PEVs have been found to enhance re-epithelialization and collagen synthesis in accelerating wound closure [25, 31]. Recently, PEVs have demonstrated potential in aiding the regeneration of corneal endothelial cells [28]. Additionally, PEVs can also contain proangiogenic factors such as TGF-β, PDGF and VEGF, which may facilitate angiogenesis. Thus, the clinical application of PEV as a delivery vehicle for the treatment of CoNV still needs further exploration.

Kaempferol (KM), a tetrahydroxyflavone, is a natural flavonoid found in a variety of plants useful to humans such as tea (family *Theacaeae*), broccoli (*Brassicaceae*), grapefruit (*Rutaceae*), and beans (*Fabacaeae*). It possesses wide pharmacological properties, including anti-angiogenic, anti-inflammatory, antioxidant, anticancer, antimicrobial, antidiabetic, neuroprotective, and cardioprotective effects [32, 33]. KM has been shown to inhibit the proliferation and migration of endothelial cells by mediating the expression of proangiogenic factors [34, 35], and has been shown to suppress CoNV [36]. However, the hydrophobic nature of KM makes it unsuitable for direct use as eye drops; its hydrophobicity aids in penetrating the lipid layer of the tear film or the corneal epithelium layer, but the aqueous tear layer and corneal stromal layer can obstruct the topical transport of KM across the ocular surface [6, 37]. To address this issue, we sought to utilize PEVs as delivery vehicles for KM.

In this study, we first formulated KM-loaded PEVs (PEV-KM) and evaluated their ability to target angiogenesis activity *in vitro*. Then, we used a mouse model of alkali burn-induced CoNV to evaluate the anti-angiogenic efficacy of PEV-KM *in vivo*.

## 2. Material and Methods

### 2.1 Materials

Kaempferol, heparin, and HEPES buffer were purchased from Sigma-Aldrich (St Louis, MO, USA). Medium 199 and trypsin-EDTA were acquired from Gibco (Grand Island, NY, USA), and fetal bovine serum (FBS) from Cytiva (Piscataway, New Jersey, USA). Endothelial cell growth supplements (ECGS) were purchased from EMD Millipore (Burlington, MA, USA). Penicillin- Streptomycin, 1-ethyl-3-(3-dimethyl aminopropyl) carbodiimide hydrochloride (EDC), 5-(and-6)- Carboxytetramethyl-rhodamine-Succinimidyl Ester (5(6)-TAMRA), 4’,6-diamidino-2-phenylindole (DPAI), and protein extraction buffer were purchased from Thermo Fisher Scientific (Waltham, MA, USA). Cell Counting Kit-8 was purchased from Dojindo Laboratories (Kumamoto, Japan). Ultra- centrifugal filters, Matrigel, Live/Dead Double Staining Kit, and MILLIPLEX MAP Mouse Cytokine/Chemokine Magnetic Bead kits were purchased from Merck Millipore (Billerica, MA, USA). Dialysis tubing (Spectra/Por, 30 kDa) was purchased from Repligen (MA, USA), and CellBrite® Cytoplasmic Membrane Dyes were purchased from Biotium (Fremont, CA, USA). Zoletil 50 solutions were obtained from Virbac Animal Health (Vauvert, Nice, France). Rompun was purchased from Bayer (Leverkusen, Germany) and topical anesthesia solution was purchased from Alcon (Hünenberg, Switzerland). Quantikine® ELISA kit was purchased from R&D Systems (Minneapolis, MN, USA). Other chemicals/reagents listed were acquired from Sigma-Aldrich.

### 2.2 Preparation of kaempferol-loading platelet-derived extracellular vesicles (PEV-KM)

Human PCs were obtained from the Taipei Blood Center (Guandu, Taiwan) and collected from voluntary non-remunerated donors who provided informed consent. The experimental procedure was approved by the Joint Institutional Review Board of Taipei Medical University (TMU-JIRB No. 201504054). Platelet-derived extracellular vesicles (PEVs) were isolated by differential centrifugation steps. First, the PC was centrifuged (3000 × *g* for 30 min at 22°C and 6000 × *g* for 10 min at 25°C) to pelletize the platelets and remove cell debris [38]. The platelet-free plasma supernatant was ultracentrifuged (25000 × *g* for 90 min at 18°C) to pelletize the PEVs. Then, the purified PEVs were resuspended in a volume of pH 7.4 HEPES buffer equal to 1/100^th^ that of the starting PCs. PEVs used in this study were pooled from 10 PCs. Exosome antibody array (cat. EXORAY200B- 4/EXORAY210B-8, System Biosciences, CA, USA) was used for EV identification, including the EV markers of CD63, CD81, ALIX, FLOT1, ICAM1, EpCAM, ANXA5, and TSG101 [39, 40].

Kaempferol (KM) was loaded into the PEV suspension thereafter. First, 100 μL PEV corresponding to 4.8 × 10^11^ particles/mL, as determined by nanoparticle tracking analysis (NTA), 10 μL KM (100 mg/mL), and 890 μL phosphate buffer saline (PBS) were gently mixed and shaken for 12 hours at 37°C. The mixture was then transferred into a centrifuge tube (30 kDa) and centrifuged (3000 × *g* for 5 minutes at 20°C) to remove unloaded KM and obtain PEV-KM for further study.

### 2.3 Characterization of PEV and PEV-KM

The particle morphology was observed using a transmission electron microscope (TEM, Hitachi HT- 7700, Tokyo, Japan) at an acceleration voltage of 75.0 kV. PEV and PEV-KM were dropped on copper grids and dried at room temperature (RT) overnight before analysis. The Dynamic Light Scattering (DLS) methodology for evaluating particle size, polydispersity (PDI), and zeta potential of nanoparticles was performed using a Zetasizer (Nano ZS 90, Malvern Instruments, Malvern UK, equilibration time: 180 seconds, temperature: 25°C). The scattering light at 90° was used for measuring/calculating the particle size [41]. The particle concentration was analyzed by Nano Sight NS300 and analyzed by NTA software 3.2 (NanoSight. Malvern Instruments). The concentration of PEV was reported as particle number/mL, and the size in mean values (nm).

### 2.4 Drug Encapsulation and release property

For determining the amount of drug loaded into the PEVs, KM was quantified by mass spectrometry. PEV-KM was first mixed with methanol at a ratio of 1:9 (v/v), and then sonicated for 30 minutes by an ultrasonic cleaner (DC200H, Delta Ultrasonic Co., New Taipei City, Taiwan) at frequency: 40 kHz and output power: 200 W to disrupt the PEV membrane. The resulting suspension was centrifuged at 12000 × *g* for 10 minutes at room temperature. The concentration of KM in the supernatant was determined using a triple quadrupole mass spectrometer (TQMS, Xevo TQ-XS, Waters, Milford, MA, USA). TQMS parameters were as follows: ionization: ESI+, capillary voltage: 2.5 kV, source temperature: 150 °C, desolvation gas flow: 900 L/h, desolvation temperature: 450°C, cone gas: 150 L/h). The results were analyzed using Waters Mass Lynx 4.1 and Quan Lynx 4.1 software. The encapsulation efficiency (EE) was calculated using the formula:

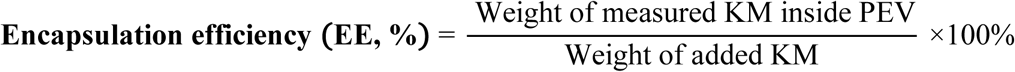

The drug release test was performed using the dialysis method. A 1 mL sample from each group was prepared and added to a dialysis bag (Spectrum™ Spectra/Por™ 2 dialysis membrane standard RC trial kit), which was then sealed. The dialysis bag was placed in a beaker containing 30 mL of release buffer (PBS with 15% ethanol) and incubated at 37 °C with stirring at 10 rpm. At specific time points (1, 2, 4, 8, 12, 24, and 48 hours) 200 μL of the buffer solution was collected and replenished with the same volume of release buffer. The concentration of KM in the collected solutions was determined using high-performance liquid chromatography (HPLC) (Hitachi L-7100, Tokyo, Japan, λ = 350 nm, a mobile phase of 20% acetonitrile, 40% methanol, 39% DI water, and 1% acetic acid) [42].

### 2.5 *In vitro* examination

Human umbilical vein endothelial cells (HUVEC) were obtained from Bioresource Collection and Research Center (BCRC, Hsinchu, Taiwan). The basal culture medium for maintaining HUVEC cells consisted of M199 medium supplemented with 10% fetal bovine serum (FBS), 25 U/mL heparin, 30 μg/mL endothelial cell growth supplement (ECGS), 1.5 g/L sodium bicarbonate, and 100 U/mL penicillin/streptomycin. The culture dishs were coated with gelatin solution (Type A) in advanced. To avoid the interference of FBS in evaluating PEV function, follow-up experiments were conducted in the same culture medium without FBS.

For the cell viability test, HUVECs were seeded in a 96-well cell culture plate at a density of 5 × 10^3^ cells/well and cultured for 12 hours at 37°C in a 5% CO_2_ atmosphere. After removing the medium, 100 μL samples in serum-free medium were added for cultivation. The concentration of PEV was set at 1% and 5% (corresponding to 4.8 ×10^9^ and 2.4 ×10^10^ particles/mL by NTA), and the KM concentration was set at 6 μg/mL and 30 μg/mL (based on the quantification of EE). Cells were co-cultured with PEV, KM, or PEV-KM as the indicated concentrations at 37°C for 24 hours. After removing the culture medium, the mixed reagent (Cell Counting Kit-8 [CCK-8]: medium = 1:10 [v/v]) was added to the wells and incubated for 3 hours. The reacted solutions were analyzed using a microplate reader (Epoch 2, BioTek, Winooski, VT, USA) at λ = 450 nm.

#### 2.5.1 Cellular uptake

To observe the cellular uptake of nanoparticles (PEV), confocal microscopy was used. HUVECs were seeded in a 96-well cell culture plate at a density of 5 ×10^3^ cells/well and cultured for 12 hours at 37°C in a 5% CO_2_ atmosphere with basal culture medium. After removing the medium, 100 μL of 1% PEV (4.8 ×10^9^ particles/mL) in serum-free medium were added and incubated at 37°C for 12 hours. Treated cells were washed twice with PBS and fixed in a 10% (w/v) formaldehyde solution. The fixed cells were stained for cell nucleus with DAPI (blue fluorescence). The KM was conjugated with red fluorescence (5(6)-TAMRA) by EDC/NHS agent for observation, and PEVs were stained with CellBrite®, a cytoplasmic membrane dye (green fluorescence). Cells were then examined using confocal laser scanning microscopy (CLSM) (TCS SP5, Leica, Solms, Germany).

#### 2.5.2 Cell migration

Cell migration was evaluated by the wound healing scratch assay. First, HUVECs were seeded in a 24-well cell culture plate at a density of 1 ×10^5^ cells/well and cultured for 12 hours at 37°C in a 5% CO_2_ atmosphere. Each well was gently cross-scratched with a 200 µL pipette tip. After removing the medium, 500 μL samples (6 μg/mL KM, 1% PEV, and 1% PEV-KM) in FBS-free medium were added and incubated at 37°C for 0, 4, and 12 hours, and observed at these times by fluorescent microscopy (IX81, Olympus, Tokyo, Japan). The wound closure area was measured by ImageJ software (1.46r, National Institutes of Health, USA) using the formula below:

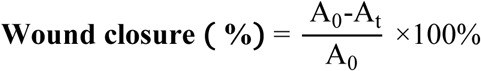

The area at time zero (A_0_) and the area after incubation time (A_t_) were used to calculate the wound closure percentage.

#### 2.5.3 Tube formation

The vessel formation capacity of HUVECs was evaluated via tube formation assay. Cells were seeded at a density of 3 ×10^4^ cells/well in a 24-well cell culture plate coated with Matrigel matrix and incubated at 37°C in a 5% CO_2_ atmosphere for 1 hour. After removing the medium, 500 μL samples (6 μg/mL KM, 1% PEV, and 1% PEV-KM) in serum-free medium were added and incubated at 37°C for 0, 4, and 12 hours, then stained with a Live Dead Cell Double Staining Kit Merck Millipore (Billerica, MA, USA). Treated cells were observed by using the fluorescent microscope (IX81, Olympus, Tokyo, Japan). The number of junctions and total tube length were measured and counted by ImageJ software (1.46r, National Institutes of Health, USA).

### 2.6 qPCR

To diminish the interference from FBS, HUVECs were adapted from 10% to 5% FBS and then to serum-free medium for gene expression examination. For the angiogenesis test, HUVECs were firstly activated by VEGF (8 ng/mL) for 6 hours, then various agents (6 µg/mL KM, PEV (1%), and PEV- KM (1% PEV with 6 µg/mL KM) in FBS-free medium were added and cocultured with HUVECs for 24 hours. The gene expression examination was conducted by real-time quantitative polymerase chain reaction (qPCR) following standard protocols of RNA extraction and reverse transcription. A StepOne Real-Time PCR System (Applied Biosystems) and the SYBR Hi-ROX Kit (Bioline, UK) were used for this examination. GAPDH was used as an internal control, and relative gene expression was quantified using the 2ΔΔCt method. The experimental condition for the angiogenic markers (VEGFA, MMP-2, and MMP-9) were tested in a 3-step cycling procedure at 95°C for 1 minute (activation), 95°C for 5 seconds (denaturation), 60°C for 10 seconds (annealing), and 72°C for 20 seconds (extension).

For evaluating the inflammatory gene expression, lipopolysaccharide (LPS, 500 ng/mL) was added to the culture medium (M199 +5%FBS) for 6 hours to induce inflammation. After removing the medium, cells were co-cultured with various tested agents (the same as aforementioned for the angiogenesis test). The parameters for the inflammation test included 2-step cycling at 95°C for 5 minutes (activation), 95°C for 1 minute (denaturation), 60°C for 1 minute (annealing), and 60°C for 10 minutes (extension). These tests specifically examined mTOR, IL-8, and MCP-1. The primer sequences are shown in **Tables S1** and **S2**. The relative gene expression levels were calculated based on the comparative 2^-ΔΔCt^ method.

### 2.7 Evaluating the PEV retention on the ocular surface

A total of 18 mice (C57BL/6J male) aged between 8 to 10 weeks were used for the *in vivo* test (n=3/group). All experimental procedures were performed following the ARVO Statement for the Use of Animals in Ophthalmic and Vision Research and approved by the Institutional Animal Care and Use Committee (IACUC) of the Taipei Medical University (IACUC approval no. LAC-2019-0140). The tested animals were first anesthetized by Zoletil/Ropum mixture (1:1,7x dilution via intraperitoneal injection). Various eye drops (KM, PEV, and PEV-KM) were carefully applied to the right eye of each mouse (5 μL/eye). The fluorescent intensity/images were analyzed/captured using an in vivo imaging system (IVIS, IVIS Lumina XRMS, PerkinElmer, Waltham, MA, USA) at 0, 1.5, 3, 4.5, 6, 7.5, 9, 15, 20, 30, 40, and 50 minutes. Native PEV was used as the control. Red fluorescent dye (5(6)-TAMRA) was conjugated with KM to observe the location of the drug, and PEVs were labelled via a membrane dye (CellBrite®) representing green fluorescence. The fluorescent content of all formulations used were adjusted to the same concentration. The fluorescence intensity of the eye, as the Region of Interest (ROI), was calculated as follows:

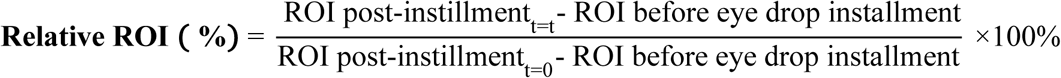

### 2.8 Evaluating the therapeutic effect of PEV-KM in a mouse model of CoNV

The animal conditions and the approval were as described above. A total of 64 mice were used in this study (n=16/group, total 4 groups). Only one eye was used for cornea NV induction, and the other eye was used as an internal control. To evaluate CoNV treatment efficiency, mice were anesthetized, and the tip of an applicator containing silver nitrate/potassium (25%/75%, Grafco, Atlanta, GA, USA) was gently pressed to the center of the cornea to induce chemical cauterization [41]. Each mouse received treatment on only one eye. 5 μL of one of the following eye drop formulations were applied to the treated eye once a day for seven days. The tested formulations included (1) PBS (negative control), (2) KM (6 μg/mL), (3) 1% PEV (4.8×10^9^ particles/mL), and (4) PEV-KM at the same PEV (1%) or KM (6 μg/mL) concentrations as their individual treating dose. The burn response and severity of corneal neovascularization were examined and imaged using a hand-held portable slit lamp (SL-17, Kowa Company, Torrance, CA, USA) in anesthetized mice. For vessel area quantification, the vessels observed from the limbus to the conjunctiva-surface layer of the sclera were manually selected and counted using ImageJ. The vessel area was calculated as follows [41]:

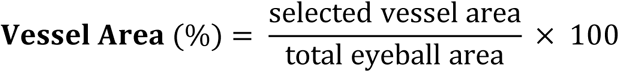

CoNV grading (from 0 to 6) was performed according to the extent of vessel growth from the limbus to the burnt edge as follows: (0) no visible vessels; (1) 1/4 distance to edge of burn; (2) 1/3 distance to edge of burn; (3) 1/2 distance to edge of burn; (4) 2/3 distance to edge of burn; (5) 3/4 distance to edge of burn; (6) vessels reach edge of burn) [41, 43]. The blister formed after alkali burn was also evaluated. A grading scale of 0 to 3 was used to identify the severity of the corneal blister: absence of a blister is graded as 0; a small blister is grade 1; a medium blister is grade 2; and a large blister formed on the central cornea is grade 3 [43].

#### 2.8.1 Immunohistological examination

For histological analysis, 2 eyeballs/group were extracted on day 7, fixed in 10% formaldehyde solution, then paraffin-embedded, and sectioned. The basic tissue architecture of the eyes was examined by hematoxylin and eosin (H&E) staining. Immunohistochemical (IHC) staining was performed as previously described [43]. Rabbit anti-mouse primary antibodies of CD31 (1:100, cat. 77699, Cell Signaling Tech., USA) and α-Smooth Muscle Actin (1:50, cat. 36110, Cell Signaling Tech., USA) were used, followed by incubation with a secondary antibody (Mouse and Rabbit Specific HRP/DAB Detection IHC kit, ab64264, Abcam Limited., USA) and hematoxylin ((HWI-125, Scytek Laboratories, UT, USA) staining. Sections were examined, and images were captured using two optical microscopes (IX81, Olympus, Tokyo, Japan; ECLIPSE Ei, Nikon, Tokyo, Japan) depending on the stain labeling.

#### 2.8.2 Protein level changes in damaged cornea

Corneas were isolated from the whole eyeball and washed twice with PBS. The collected corneas (4 eyeballs per group) were pooled together and then digested using a protein extraction buffer. The cornea lysate was further homogenized using a bead beater-type homogenizer (Beads crusher μT-12, TAITEC, Saitama, Japan). The supernatant was collected after centrifuging the mixture from each sample. Total protein was quantified by Bradford assay (p010, GeneCopoeia, Rockville, MD, USA), and adjusted to the same concentration. Quantification of angiogenic cytokines (VEGF) and inflammatory factors (TNF-α, IL-1β, IL-6, and MCP-1) in different therapeutic agents were measured by Quantikine® ELISA kit (R&D Systems Inc. Minneapolis, MN, USA) in triplicate. All experiments were conducted according to the manufacturer’s protocols, with 3 independent biological repeats for each group.

### 2.9 Statistical analysis

All results are presented as mean ± standard deviation (SD). Each group was tested with sample sizes ranging from 3 to 6. For statistical analysis, one-way analysis of variance (ANOVA) was used to compare multiple groups, performed using SPSS 18.0 (SPSS, Chicago, IL, USA) or GraphPad Software (version 8.0; GraphPad Software, USA). A p-value < 0.05 was considered to be statistically significant.

## 3. Results

### 3.1 Characterization of PEV and PEV-KM

We first isolated and confirmed EV the purified EVs were released from platelets. The presence of EV markers such as CD63, ANXA5, TSG101, ALIX, CD81, and FLOT1, as well as cell adhesion protein markers ICAM and EpCAM, and light expression of Golgi marker GM130 indicated that the isolated particles were EVs (**Fig. S1**). Following the optimal parameters of PEV from our previous study [38], the dynamic light scattering (DLS) results revealed that PEVs with a diameter of 144.3 nm, and ζ potential at -5.8 mV were acquired (**Table 1**).

**Table 1.** Characterization of PEV and PEV-KM.

| Sample | Size<br>(nm) | PDI<sup>1</sup> | $\zeta$ potential<br>(mV) | E.E.<sup>2</sup><br>(%) | Concentration<br>(number/mL) | Recovery<br>rate (%) |
| --- | --- | --- | --- | --- | --- | --- |
| PEV | 144.3 ± 19.2 | 0.46 ± 0.03 | -5.8 ± 1.5 | / | 4.8×10<sup>11</sup> ± 3.2×10<sup>10</sup> | / |
| PEV-KM | 161.5 ± 29.5 | 0.41 ± 0.04 | -13.7 ± 0.8 | 61.0 ± 5.2 | 4.5×10<sup>11</sup> ± 3.6×10<sup>10</sup> | 94.9 ± 3.2 |

<sup>1</sup> PDI: Polymer dispersity index. <sup>2</sup> E.E.: Encapsulation efficiency of KM. n=3~5.

After loading with KM, the diameter of PEV-KM slightly increased to 161.5 nm; and the ζ potential became more negative (-13.7 mV). The decreased zeta potential may result from the addition of KM, which has a hydroxyl group (OH^-^), indicating the successful loading of KM in PEV. The PEV and PEV-KM had a spherical shape with a diameter < 200 nm, and no particle aggregation was observed by Transmission Electron Microscopy (TEM; **Fig. 1A)**. The enlarged images of PEV revealed a membrane-like structure covering the particles, and KM-loaded PEV forms granules inside the particle were observed (**Fig. 1A**).

**Figure 1.**
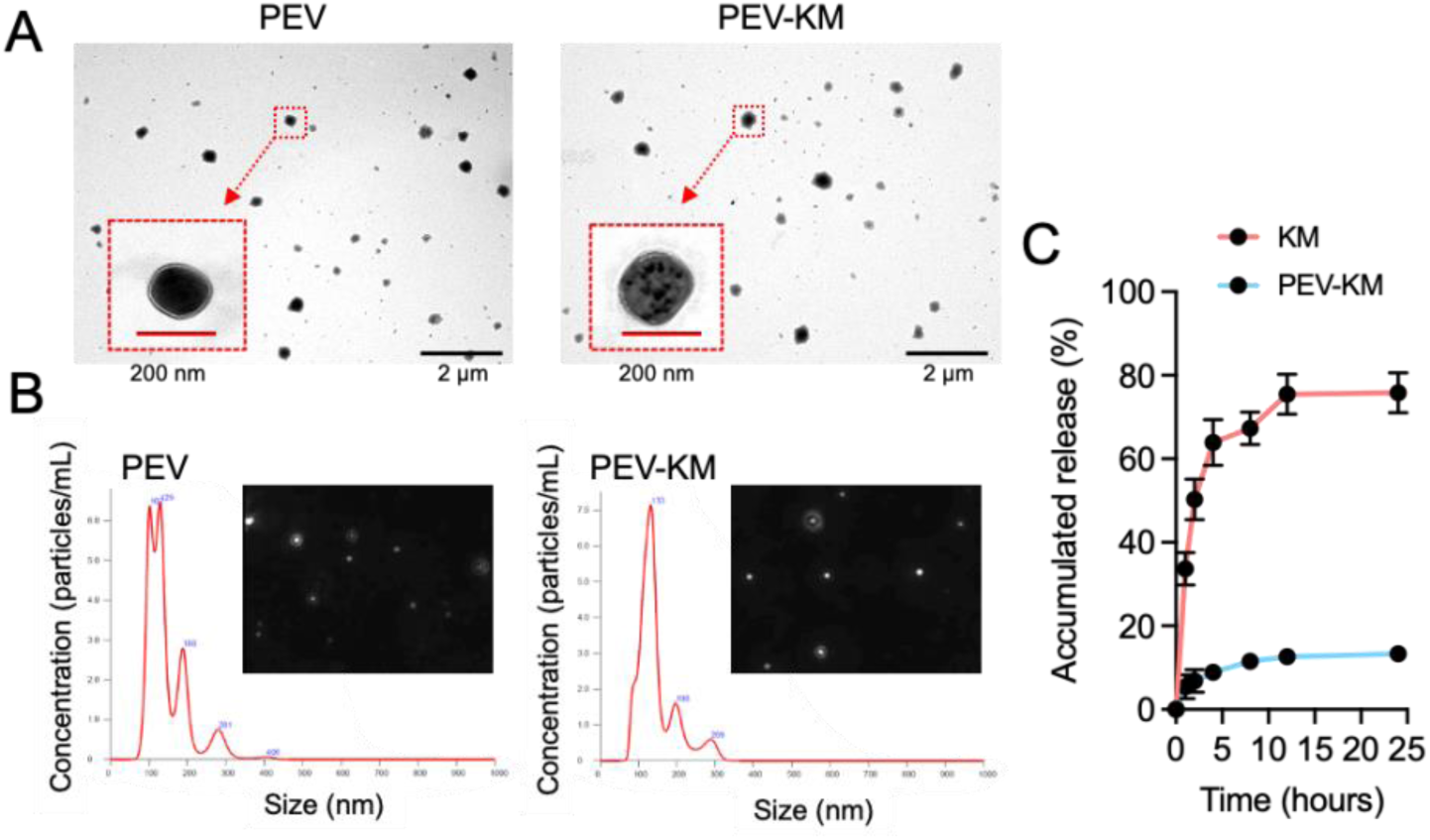
Characterization of PEV and PEV-KM. (A) TEM images of PEV and PEV-KM. (B) NTA results of PEV and PEV-KM. (C) The drug release profile of KM and PEV-KM was evaluated in PBS (pH 7) with a 15% ethanol addition.

In **Table 1**, the PDI of PEV and PEV-KM were ∼0.4, indicating a mono-dispersed condition. This is also supported by the results of NTA examinations (**Fig. 1B**), as the majority of particle sizes ranged from 100 to 200 nm. PEV and PEV-KM still exhibit Brownian motion as a colloidal solution. In addition, NTA results showed that the concentration of PEV was approximately 4.8x10^11^ particles/mL (**Table 1**). After KM loading, a centrifugation process for separating unloaded KM had minimal impact, and the number of PEV-KM was maintained at 4.5×10^11^ ± 3.6×10^10^ particles/mL (**Table 1**). The recovery rate of PEV-KM was high (94.9%, **Table 1**).

The concentration of KM within the PEV was determined by TQMS by directly measuring KM from the dissociated PEV. The mean encapsulation efficiency (EE%) of PEV-KM was found to be 61.0 ± 5.2 (**Table 1**). Due to the poor water solubility of KM, the release experiment was conducted in PBS containing 15% ethanol. The release profiles are shown in **Fig. 1C**. After 24 hours, 75.8% of KM had diffused outside the dialysis bag in the KM group, whereas only 13.3% of KM from the PEV- KM was released from the nanocarrier, PEV. Our data suggest that PEV-KM has a slow-release pattern.

### 3.2 *In vitro* cellular uptake and cell viability

To confirm the localization of nanocarriers (PEV) and the drug (KM) in cells, confocal laser scanning microscopy (CLSM) was used to observe the fluorescence-labelled PEV-KM intracellularly. The results are shown in **Fig. 2A**. Both the red fluorescent signals (indicates KM) and the green fluorescent (indicates PEV) were found in the cytoplasm and distributed around the nucleus. Quantification of the KM content in cells via fluorescent intensity is revealed in **Fig. 2B**.

**Figure 2.**
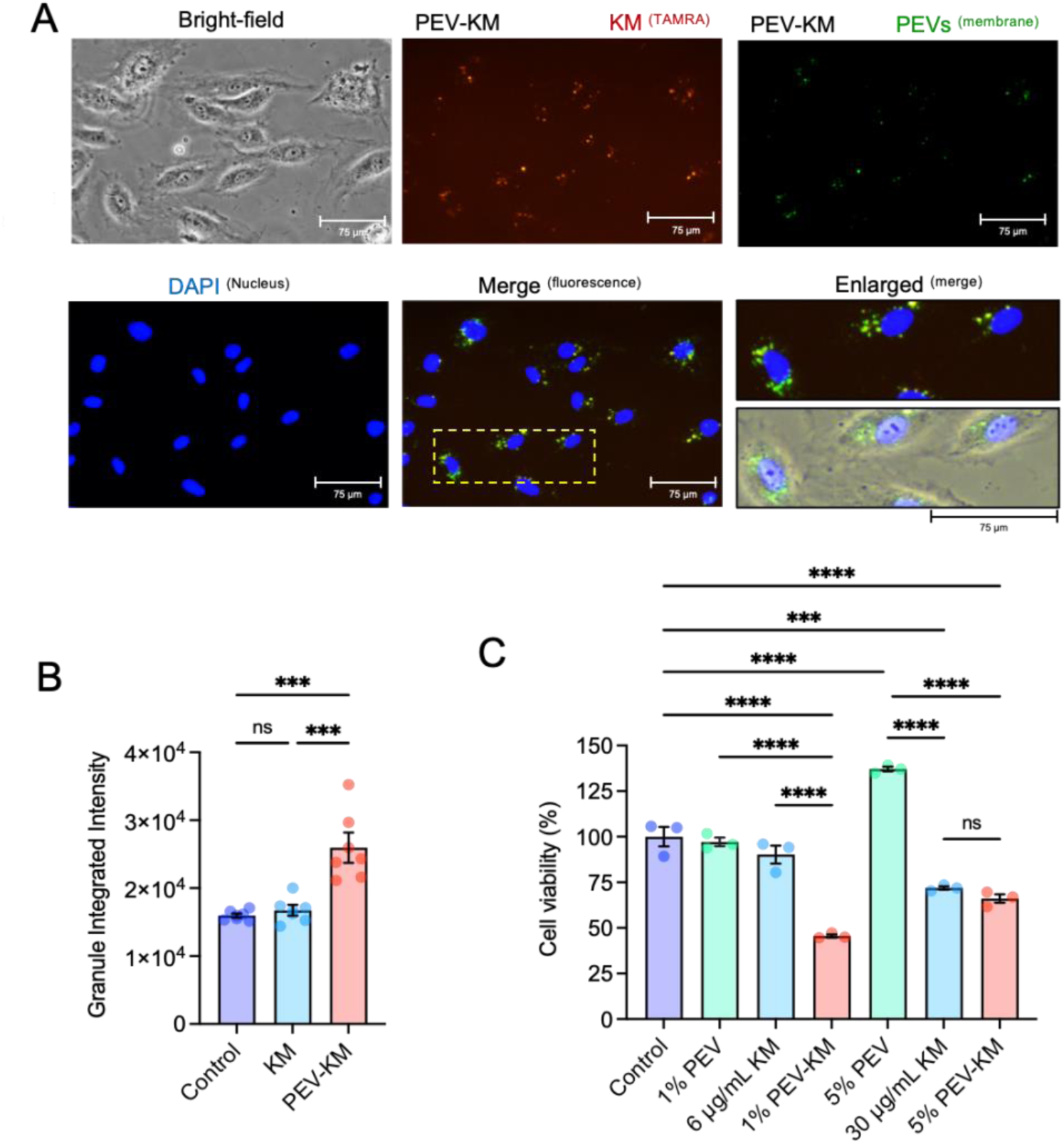
In vitro cellular uptake and cytotoxicity. (A) Images acquired from HUVEC treated with PEV-KM for 12 hours. PEVs were labelled in green fluorescence, KM by red fluorescence (TAMRA), and the nucleus is shown in blue (DAPI). (B) Quantitation of the KM content in cells via fluorescent intensity (n = 6-7). (C) Cell viability was tested by the WST1 agent after 24 hours of incubation with variant PEV or KM formula (n = 3). Data presented as mean ± SEM. Statistical analysis was conducted using one-way ANOVA and Tukey’s multiple comparison test (B, C). ns: not significant, ***p ≤ 0.001, ****p ≤ 0.0001.

The mean fluorescence intensity in PEV-KM treated cells (25,963) was significantly higher than in KM-treated cells (16,736, n = 6-7; p < 0.001), indicating that a significantly higher amount of KM can be transported into cells when loaded within the PEV vehicles. To investigate the effect of PEV or KM on cells, the cell viability of HUVECs treated with PEV, KM, and PEV-KM was assessed using a Cell Counting Kit-8 (CCK8) assay. The cell viability of the 1% PEV-treated cells was 97.1% ± 2.4% and then increased to 137.1% ± 1.3% when treated with 5% PEV compared to control cells (100% ± 5.4%), indicating greater metabolic activity or proliferation in a high PEV concentration. The cell viability of the KM-treated cell was 90.2% ± 4.9% and 71.9% ± 0.9% at KM concentrations of 6 μg/mL and 30 μg/mL, respectively, compared to control cells. Our result shows that a higher dose of KM may cause toxicity or affect cellular metabolic activity. Interestingly, a reduction in the cell viability was found in both the 1% PEV-KM (45.6% ± 1.2%, n = 3; p < 0.0001) and 5% PEV-KM (66.1 ± 2.3%, n = 3; p < 0.0001) compared to control cells (**Fig. 2C**). This result was attributed to the capacity of PEV to enhance KM internalization by HUVEC and increase the toxic effect compared to KM alone.

### 3.3 PEV up-regulates, but PEV-KM down-regulates angiogenic and inflammatory genes *in vitro*

Previous studies indicate that PEV is activated in response to angiogenesis and aids in the suppression of inflammation. We first assessed the effect of PEV, KM and PEV-KM on angiogenesis-related gene expression upon VEGFA stimulation in human endothelial cells (HUVECs). Consistent with the literature, PEV treatment clearly induced the upregulation of the angiogenesis-associated genes such as VEGFA, MMP2 and MMP9 (**Fig. 3A**). Interestingly, we found that the effects were attenuated when KM was loaded into PEV (PEV-KM) (**Fig. 3A**). These results suggest that PEV alone has a strong effect on the stimulation of angiogenesis. However, this effect can be managed by loading an anti-angiogenic agent such as KM to suppress pathological angiogenesis.

**Figure 3.**
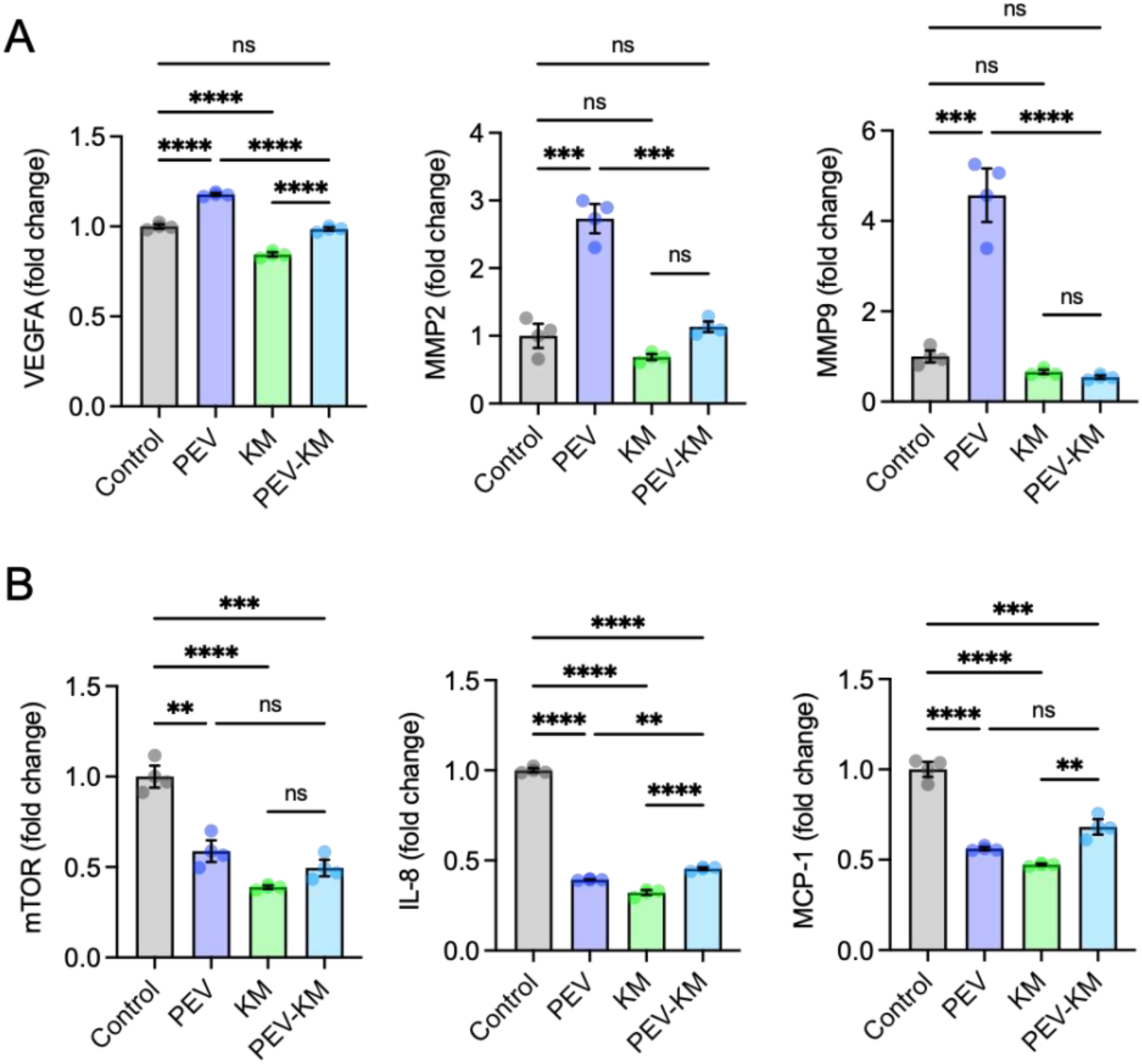
The effect of PEV, KM or PEV-KM formula on angiogenesis and inflammation-associated gene expression. (A) The effect of angiogenesis-associated gene expression (VEGFA, MMP2, and MMP9) by various PEV formulations under stimulation with VEGFA (n=3). (B) The effect of inflammatory-associated gene expression (mTOR, IL-8, and MCP-1) by various PEV formulations under stimulation with VEGFA (n=3). Data presented as mean ± SEM. Statistical analysis was conducted by one-way ANOVA and Tukey’s multiple comparison test (A and B). *p ≤ 0.05, **p ≤ 0.01, ***p ≤ 0.001, ****p ≤ 0.0001.

We further investigated the role of PEV and PEV-KM in response to inflammatory insults to HUVECs. Our results showed that PEV treatment alone effectively inhibited the expression of LPS- induced proinflammatory genes, including mTOR, IL8, and MCP-1 (**Fig. 3B**). A similar effect was also found in KM or PEV-KM treated cells (**Fig. 3B**). Collectively, our data confirmed the utilization of PEV-encapsulated KM for mediating angiogenesis and inflammation response in human endothelial cells.

### 3.4 Effects of PEV and PEV-KM on endothelial cell activity

Endothelial cell migration and tube formation assays were used in this study to evaluate the effect of PEV and PEV-KM on endothelial cell activity. For the cell migration assay, the wound closure of human umbilical vein endothelial cells (HUVECs) treated with 1% PEV, 6 μg/mL KM, or 1% PEV- KM were examined. Changes in the wound area and quantitative data are shown in **Fig. 4**.

**Figure 4.**
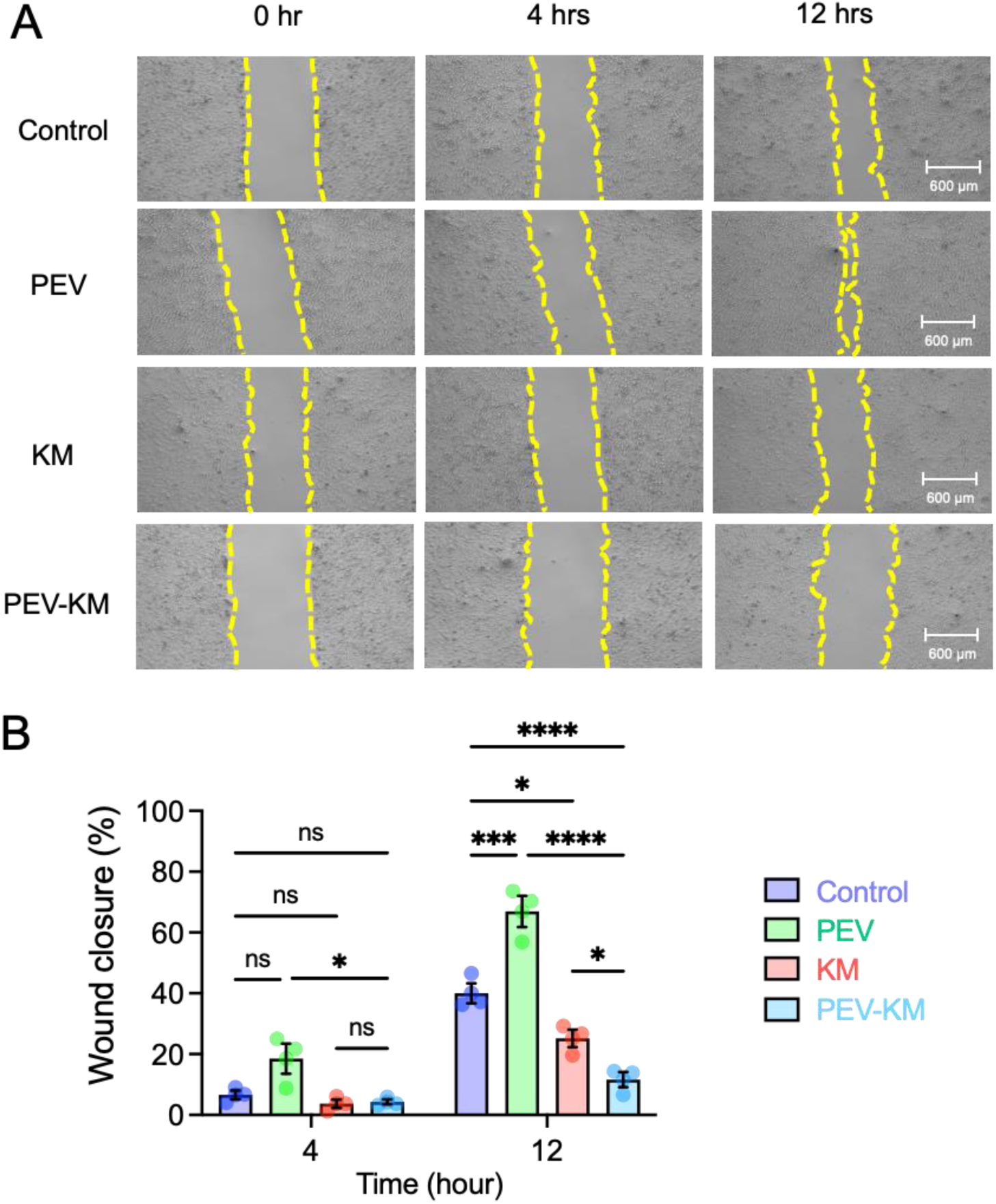
The effect of PEV, KM or PEV-KM formula on endothelial cell migration. (A) Images of the wound area closure were taken after scraping 0, 4, and 12 hours. (B) Quantification of wound closure at 4 and 12 hours, determined by the ratio of cells in the gap area (n = 3). PEV: 1% PEV, KM: 6 mg/mL KM, and PEV-KM: 1% PEV-KM. Data presented as mean ± SEM. Statistical analysis was conducted using two-way ANOVA and Tukey’s multiple comparison test (B). ns: not significant, *p ≤ 0.05, ***p ≤ 0.001, ****p ≤ 0.0001.

The wound closure area in control (6.6% ± 1.4%), KM-treated (3.7% ± 1.4%), and PEV-KM- treated (4.3% ± 0.8%) groups was similar, with wound closure ratios ranging from 3.7% to 6.6% at the 4-hour timepoint. However, the PEV-treated group (18.5% ± 4.9%) had a slightly higher closure ratio compared to the other 3 treatment groups. At the 12-hour timepoint, a narrowed gap area was observed in the control group where the closure ratio increased to 40% ± 3.3%. In contrast, the PEV-treated group showed a superior wound closure ratio (66.9% ± 5.1%) compared to the control group (p < 0.0001). However, a significant reduction in cell migration was observed in both KM-treated (25.2% ± 2.9%; p < 0.05) and PEV-KM-treated (11.6% ± 2.5%; p < 0.0001) groups at the 12-hour timepoint compared to the control group (**Fig. 4**). Together, our data suggest that PEV promotes endothelial cell migration, whereas KM and PEV-KM inhibit the process. The results of the tube formation assay are shown in **Fig. 5**.

**Figure 5.**
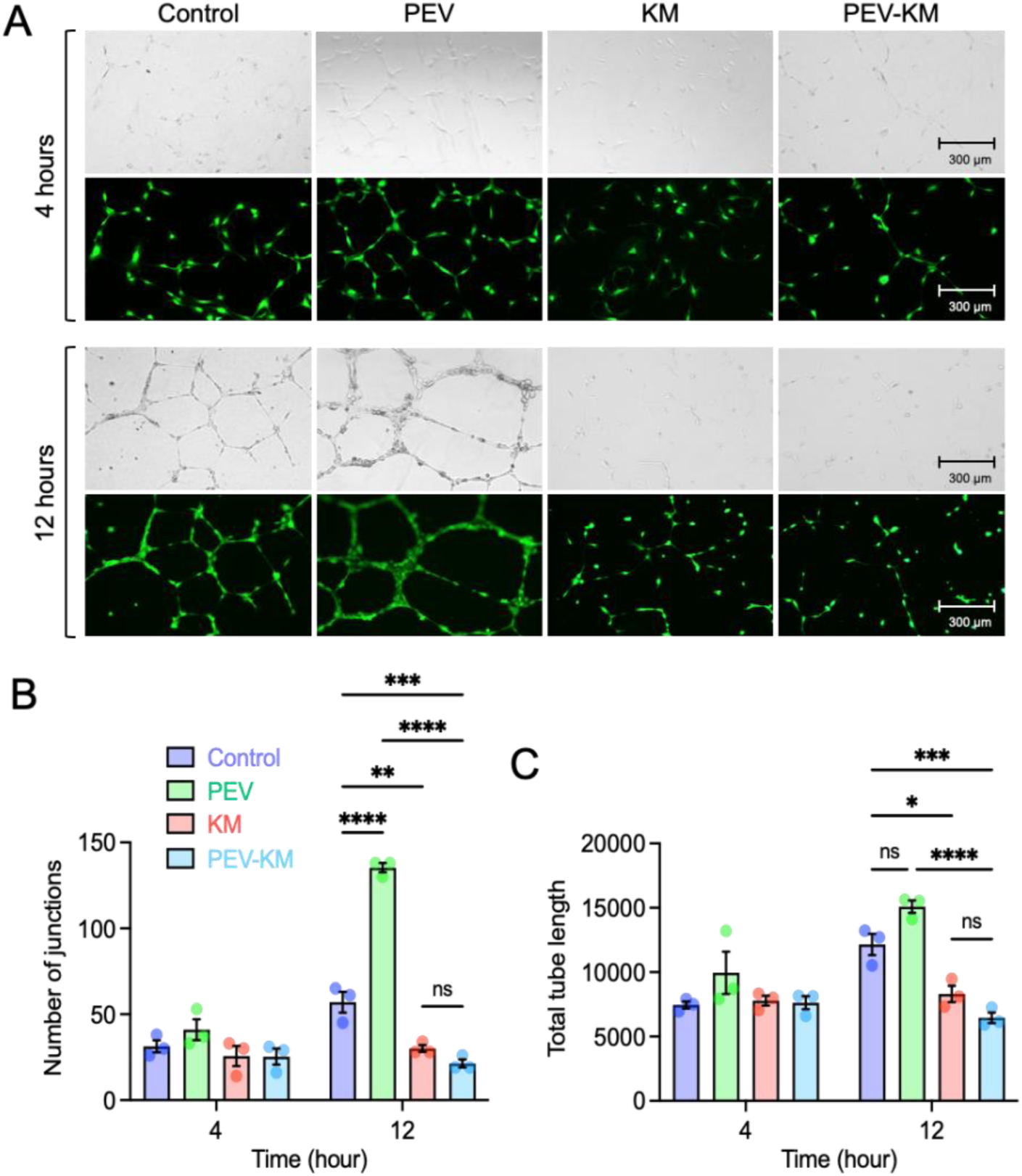
The effect of PEV, KM or PEV-KM formula on endothelial tube formation capability. (A) Images of tubes under bright field and fluorescence examination taken after 4 and 12 hours of cultivation. (B) The number of junctions counted at 4 and 12 hours. (C) Total tube length calculated at 4 and 12 hours. Data presented as mean ± SEM. Statistical analysis was conducted using two-way ANOVA and Tukey’s multiple comparison test (B, C). ns: not significant, *p ≤ 0.05, **p ≤ 0.01, ***p ≤ 0.001, ****p ≤ 0.0001.

In the control group, HUVECs established a partial net-like structure at the 4-hour timepoint and preserved this network for up to 12 hours (**Fig. 5A**). Similar to the cell migration assay, the 1% PEV-treated group exhibited a well-formed tube network as early as 4 hours, and network integration was found after 12 hours. In contrast, cells treated with KM or PEV-KM showed no tube formation and displayed random cell distribution on matrigel (**Fig. 5A**). We then quantified the junctions and tube lengths of the vessel network, as shown in **Fig. 5B and 5C**. At the 4-hour timepoint, there was no statistical difference between each treatment group compared to the control group. However, an increase in the junctions (135 ± 3; p < 0.0001) and tube lengths (15093 ± 484; p=0.0655) was observed in the PEV-treated group compared to the control group (junctions: 67 ± 6; tube lengths: 12152 ± 831) at the 12-hour timepoint (**Fig. 5B and 5C**). Moreover, an inhibitory effect of KM and PEV-KM was observed in both the junctions (KM: 30 ± 2, p < 0.01; PEV-KM: 64 ± 4, p < 0.001) and tube lengths (KM: 8304 ± 629, p < 0.05; PEV-KM: 6454 ± 409, p < 0.001) compared to the control group (**Fig. 5B and 5C**). Overall, our data indicate that PEV-KM is the most effective inhibitor of tubular formation in endothelial cells.

### 3.5 PEV enhancing the drug retention on the ocular surface of mouse cornea

To confirm that the ocular retention of the drug can be improved by encapsulating it inside PEV, PEV and KM were labelled with a membrane dye (CellBrite®, green fluorescence) and TAMRA (red fluorescence) and topically applied to mouse eyes. The fluorescent signals (ROI) on the ocular surface from each treatment group were examined using IVIS. A decrease in the fluorescence signals were observed for both the PEV-CellBrite® and PEV-KM-CellBrite® treated mice, dropping from 100% to 72% within the first 10 minutes, and decreasing to 67.3% by 50 minutes (**Fig. 6A**). This indicates that KM loaded in PEV (PEV-KM) does not affect PEV properties as an effective nanocarrier for drug delivery to the ocular surface (**Fig. 6B**).

**Figure 6.**
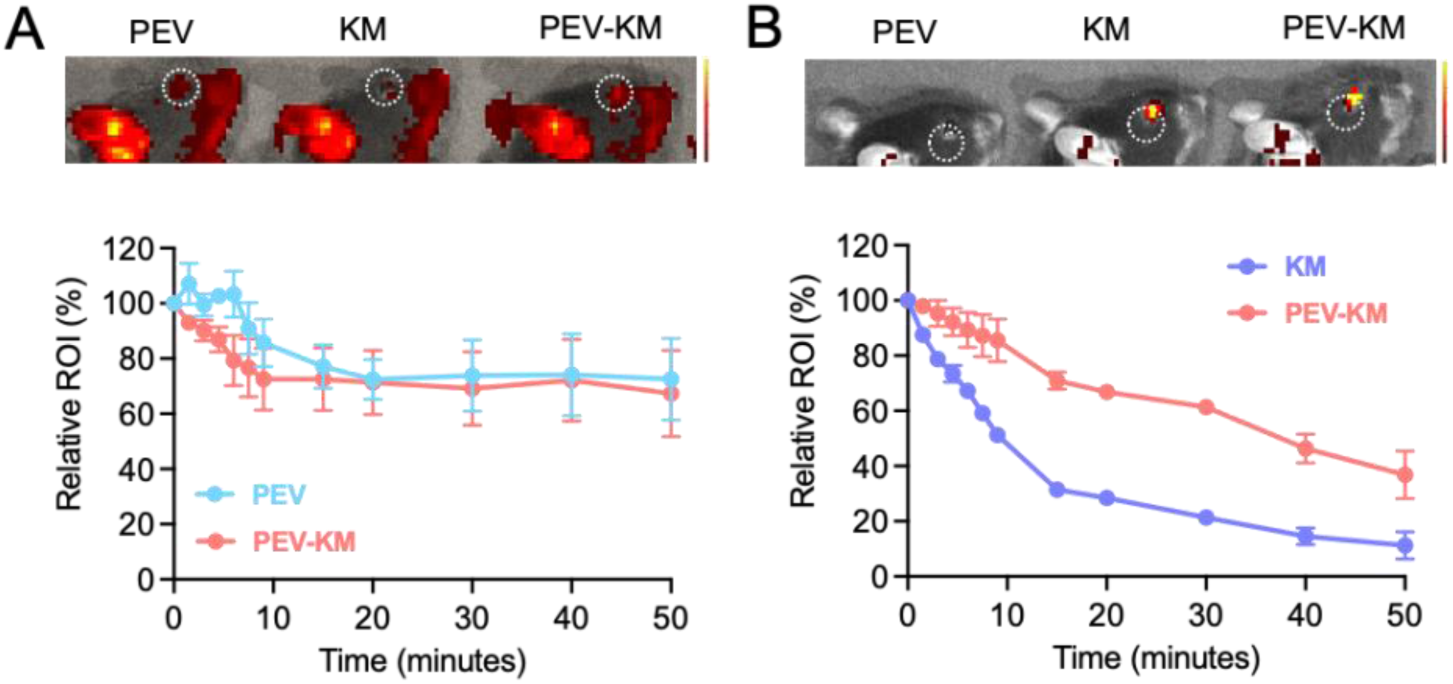
PEV-encapsulated KM is retained in the mouse cornea. (A) A fluorescent signal was acquired from PEV with green fluorescence labelling on its cytoplasmic membrane after receiving eyedrops for 50 minutes, and the ROI quantification was acquired at variant time points. (B) Red fluorescence (TAMRA) representing KM as the fluorescent signal after dosing for 50 minutes and its quantification results. The percentage change in fluorescent intensity (ROI) was individually compared with the initial ROI. Data presented as mean ± SEM.

A notable difference in the retention timeline of KM-TAMRA and PEV-KM-TAMRA in mouse eyes was observed. After 10 minutes of dosing, the fluorescent signals of KM-treated mice dramatically reduced to 30% and continued to decrease to 11.2% by 50 minutes. The relative fluorescent signals of PEV-KM-treated mice remained around 80% after 10 minutes, then gradually reduced to 50% at 50 minutes, which was significantly higher than the KM-treated mice at all time points. Our data further indicate that PEVs are able to sufficiently retain KM on the ocular surface.

### 3.6 Topical application of PEV-KM relieves corneal neovascularization in the mouse cornea

An alkali-burned corneal injury was used to induce CoNV in the mouse eye. The experimental timeline is shown in **Fig.7**. Daily dosing of variant eyedrop formulations were tested, including PBS, 1% PEV, 6 μg/mL KM, and 1% PEV-KM. A clear and transparent cornea was observed in the normal eye. The radial growth of blood vessels from the corneal limbus towards the central burn area was observed 7 days after chemical cauterization (**Fig. 7B**). Neovascularization grading showed that PBS-treated mice had the highest score of 6 (7/12 score of 6), suggesting that severe neovascularization had occurred in the cornea. The score was significantly reduced in topical PEV (7/12 score of 3-4), KM (9/12 score of 2-3) and PEV-KM (12/12 score of 0-1) treated mice (**Fig. 7C**). The corneal blister grading showed a similar trend to the neovascularization grading. The PBS-treated mice had the most severe blister score (10/12 score 1-2). The score was significantly reduced in topical PEV-KM (11/12 score of 0-1) treated mice (**Fig. 7D**).

**Figure 7.**
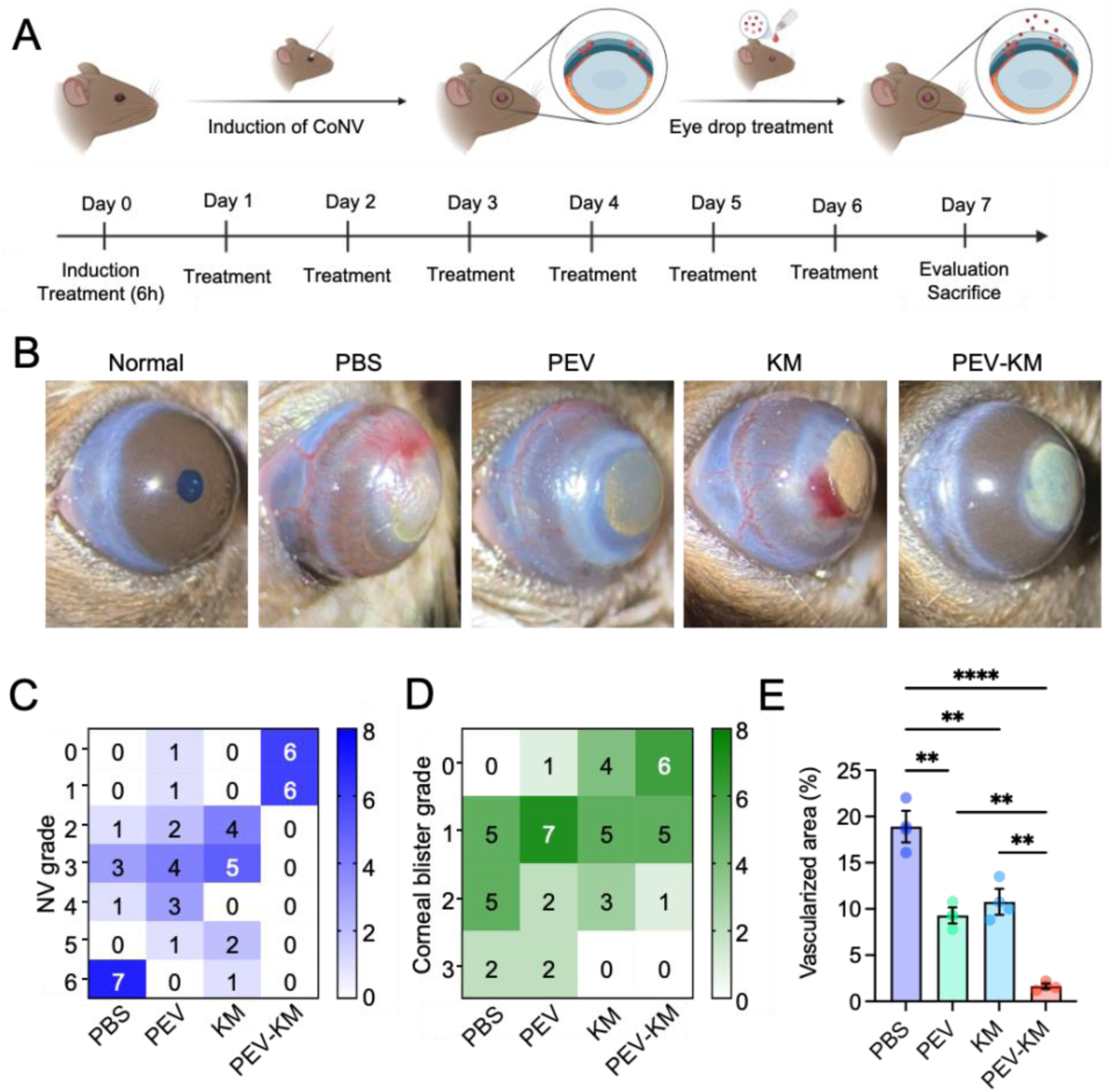
The effect of PEV, KM and PEV-KM treatment in a mouse model of CoNV. (A) Schematic diagram of the timeline for CoNV treatment with eyedrops. C57BL/6J mouse eyes were chemically cauterized following a daily eyedrop application of PBS, GNPs, PEV, KM or PEV-KM for 7 days. Created with BioRender.com. (B) Representative images were taken 7 days post-treatment. (C) Quantification of neovascularization (NV) grade according to the extent of CoNV was determined by the extent of vessel growth from limbus to burn edge (refer to methods for grading protocol) (n = 12 eyes per treatment group). (C) Quantification of corneal blister grade according to the severity of the corneal trauma/burn was determined by the fluid filling a space between layers of corneal tissue (refer to methods for grading protocol) (n = 12 eyes per treatment group). (D) Quantification of vessel area (normalized to total cornea area) in the cornea (n = 3 corneas per treatment group). Data presented as mean ± SEM. Statistical analysis was conducted by one-way ANOVA and Tukey’s multiple comparison test (E); **p ≤ 0.01, ****p ≤ 0.0001.

Quantitative analysis of CoNV revealed a significant reduction of neovascular area in mice treated with PEV, KM, PEV-KM compared with PBS-treated mice (**Fig. 7E**). The PBS-treated group showed abundant vessel formation from the limbus to the central cornea, with the largest vascularized area (18.9 ± 1.7%). After treatment with PEV or KM, the vascularized areas in both groups were significantly reduced (PEV: 9.3 ± 0.9%, p < 0.01; and KM: 10.8 ± 1.4%, p < 0.01) compared to the PBS-treated mice. The PEV-KM-treated mice demonstrated the lowest vascularized area (1.6 ± 0.3%, p < 0.0001, compared to the PBS-treated mice), with almost no vessels observed on the cornea, suggesting that PEV-KM provided a superior anti-angiogenic effect.

### 3.7 Topical application of PEV-KM reduces inflammatory cytokines in the mouse cornea

Corneal vasculatures and scarring/fibrosis were further examined by immunohistochemistry analysis. In the PBS, PEV and KM-treated eyes, there was a significant increase in inflammatory cell infiltration (**Fig. 8A, H&E**; green arrows) and vascular lumens (**Fig. 8A**, CD31; red arrows) in the corneal stroma. There is less evidence of corneal inflammatory and angiogenic response in the PEV- KM-treated eyes (**Fig. 8A**). The expression of alpha-smooth muscle actin (α-SMA) in the corneal stroma reflects the development of corneal scarring and fibrosis. The eyes treated with PBS and KM exhibited strong α-SMA positive staining in the stroma, indicative of corneal fibrosis whilst both the PEV- and PEV-KM-treated eyes had less α-SMA staining, suggesting that PEV may have a role in mediating the fibrotic response in corneal tissue following chemical cauterization (**Fig. 8A**, α-SMA).

**Figure 8.**
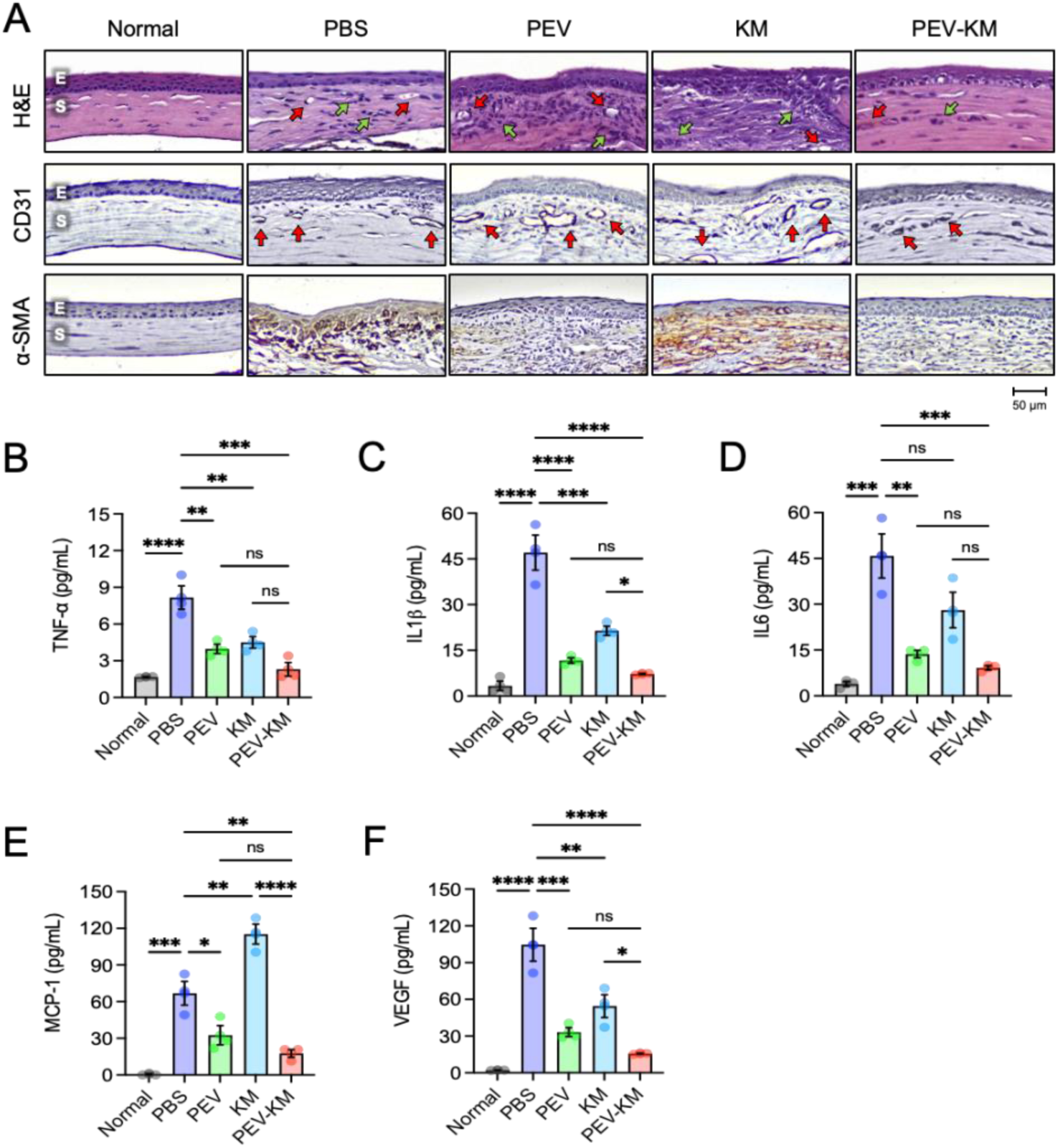
PEV-KM suppresses CoNV and reduces inflammatory cytokines in the mouse cornea. (A and B) Histological examination of mice cornea. All the tissues were acquired on day 7. (A) H&E staining and immunohistochemical staining of CD31 and α-SMA. Scale= 50 μm. E: epithelium, S: stroma. Red arrows: neovascularization, green arrows: inflammatory cell infiltration. (B-F) The protein level of inflammatory cytokines (TNF-α, IL-1β, IL-6, MCP-1, VEGF and MMP2) from corneal lysate (n = 3 corneas per treatment group) was examined by ELISA. Data presented as mean ± SEM. Statistical analysis was conducted by one-way ANOVA and Tukey’s multiple comparison test (B-F). *p ≤ 0.05, **p ≤ 0.01, ***p ≤ 0.001, ****p ≤ 0.0001.

Numerous studies have reported that various pro-inflammatory cytokines play critical roles in the pathogenesis of CoNV [6, 44–46]. Consistent with these findings, we found that the increase of pro-inflammatory cytokines such as TNFα, IL-1β, IL-6 and MCP-1 protein was significantly increased in the mouse model of CoNV after 7 days of chemical cauterization (**Fig. 8B-E**, PBS-treated eyes; p < 0.001), suggesting relatively chronic inflammatory responses were developed in this model.

Interestingly, we found that pro-inflammatory cytokines were decreased in the eyes treated with PEV, while the protein expression (TNFα and IL-1β) was moderately reduced in those eyes treated with KM (**Fig. 8B-E**). Moreover, these pro-inflammatory cytokines were further reduced in PEV-KM-treated eyes. Similar results were observed in the expression of pro-angiogenic protein VEGF (**Fig. 8F**). Together, our data indicate that PEV-KM efficiently suppresses inflammatory responses and inhibits pathological angiogenesis in the cornea in chemical cauterization-induced CoNV.

## 4. Discussion

We sought to design a novel therapeutic strategy for CoNV as current treatments may not be effective or suitable for some patients. For example, bevacizumab is an approved treatment for CoNV that can be applied topically or injected into the subconjunctival space. However, topical administration of bevacizumab can increase the risk of corneal epithelial defects over time, and CoNV may recur after successful treatment with subconjunctival injections, meaning follow-up injections are required [47, 48]. Topical steroids have also been widely used to treat CoNV but can not fully reverse established CoNV and are associated with side effects like accelerated cataract formation, steroid-induced glaucoma, and local immunosuppression [49]. In the present study, we investigated the utility of PEVs for delivering the anti-angiogenic agent kaempferol (KM) in an eye drop formulation for the treatment of CoNV. Our data indicate that the PEV-KM formulation outperforms KM-only administration in suppressing CoNV in a mouse model. This superior performance is due to the combined effects of PEVs’ anti-inflammatory properties and the prolonged retention of KM on the ocular surface. Thus, the PEV-KM formulation presented here may offer an alternative to existing therapies, offering enhanced therapeutic efficacy with potentially fewer adverse effects.

PEVs hold considerable promise as an innovative drug delivery platform for CoNV. They are the most abundant EVs in the blood and are relatively easy to obtain and use in large quantities, allowing the possibility of scale and clinical translation [20, 50]. Additionally, PEVs inherently possess therapeutic properties due to their content of various bioactive molecules, such as growth factors, antioxidants, genetic material and lipids [24]. The therapeutic potential of PEVs as a drug delivery platform has been demonstrated in other physiological systems. In 2021, Ma et al. described the delivery of dexamethasone combined with PEVs to mice with pneumonia, characterised by cytokine storms and inflammation in the lung [51]. PEVs carrying only 0.25× of the normal dexamethasone dose were able to relieve the cytokine storm in mice. This result is suggestive that PEVs are effective at delivering drugs in a targeted manner. The same group also evaluated PEVs for rheumatoid arthritis (RA), loaded with plant-derived berberine (BBR). Their BBR-PEV formulation was able to reduce ankle (hindpaw) swelling by 18.4%, and the authors found that PEV accumulated in the damaged joint, suggesting that PEVs have chemoattractive potential [52].

There are several approaches to loading drugs into EVs, including incubation, sonication, freeze- thaw, saponification, and electroporation [17, 53]. Among these, incubation is relatively simple and does not affect the EV’s structure, although it may result in lower drug encapsulation efficiency (E.E.). This can be improved by adding membrane permeabilizer/permeability enhancers [53, 54]. In this study, PEV-KM was prepared by dissolving KM in dimethyl sulfoxide (DMSO), a common permeability enhancer, followed by incubation with PEV. The encapsulation efficiency of KM in PEVs exceeded 60%, without compromising the structural integrity of the vesicles. Similar to EVs, liposomes – characterized by a phospholipid bilayer membrane – have been confirmed to incorporate hydrophobic KM inside the hydrophobic zone of liposomes according to the partition coefficient [55]. This result supported our hypothesis that KM was distributed inside the hydrophobic bilayer of PEVs. Fluorescently labelled PEVs were found to be readily internalized by cells, accumulating in the perinuclear region, indicating that PEV was indeed taken up into HUVEC through endocytosis. This suggests that PEV-KM, as a nanomedicine, can be internalized through EV-mediated endocytosis, allowing for the gradual intracellular release of KM to exert its therapeutic effects.

Poor drug bioavailability (<5%) of eyedrops is attributed to short retention in eyes due to ocular barriers [6, 37, 56], making effective drug delivery to the eye still a challenge. Advantages of nanoparticles include prolonged retention on the ocular surface, protection of drugs from degradation, sustained drug release, and a reduction in both side effects and dosing frequency [11, 12, 37, 57]. Among various kinds of artificial nanoparticle systems (chitosan, gelatin, liposome etc,), natural nanoparticles, such as EVs, have recently gained high-profile attention for the mentioned advantages and their remarkable ability to cross natural tissue barriers. The corneal epithelium and the stroma are the main barriers in the cornea that limit topical drug delivery [20, 58]. EVs can cross various biological barriers, including tissue barriers, cellular barriers (plasma membranes), and intracellular barriers (across endosomal membranes) [59]. EVs have been shown to cross the blood-brain barrier (BBB) [59, 60]. For example, intravenous injection of RVG-exosomes loaded with siRNA for delivery to the brain for Parkinson’s disease (PD) treatment has been used successfully *in vivo* to reduce intraneuronal protein aggregates via enhanced exosome cargo passing through the BBB [59, 60].. Wu et al. examined and proved that cell uptake of PEVs primarily involved endocytosis, which occurred via a clathrin-dependent pathway [38]. Heusermann et al. demonstrated EVs can be internalized more efficiently than synthetic nanocarriers such as lipid nanoparticles; EVs entered cells within minutes of addition, while lipid NPs accumulated at the cell surface first [61, 62]. From the uptake and ocular retention results, PEV or PEV-KM could overcome cellular barriers to be transferred into cells and exhibit higher/longer retention on the ocular surface via crossing corneal barriers. The ability of nanocargo such as PEV to help retention on eyes to increase drug content was proved and is consistent with our previous study on gelatin-, chitosan-made nanoparticles [42, 63]. Our analysis of PEV-KM release profiles over a 24-hour period shows that coupling of KM within a PEV delivery system allows for the slower release of KM, potentially prolonging the drug’s residence time within the eye and thereby extending its therapeutic effect. Based on our uptake and ocular retention studies, PEV or PEV-KM could overcome cellular barriers to be transferred into cells and exhibit higher/longer retention on the ocular surface by crossing corneal barriers. The ability of nanocargo such as PEVs to aid ocular retention to increase drug delivery is proven and is consistent with our previous study on gelatin and chitosan-made nanoparticles [42, 63].

PEVs offer significant promise beyond their nanoscale size. They exhibit both active and passive immunomodulatory effects, with notably reduced immunogenicity, suggesting future applications in allotransplantation and personalized medicine [64]. *In vitro*, PEVs alone were able to inhibit the expression of proinflammatory genes, including mTOR, IL8 and MCP-1, in human endothelial cells exposed to inflammatory stimuli. Similarly, both KM-only and PEV-KM treated cells demonstrated anti-inflammatory effects, consistent with the well-documented anti-inflammatory properties of KM [32]. Apart from active immune targeting, PEVs also express various other proteins that assist in actively targeting damaged vessels and may be responsible for the angiogenic effect we found in the present study. This aligns with PEVs’ known ability to promote angiogenesis, likely due to their rich content of growth factors, including VEGF, PDGF, IGF-1, EGF, and FGF [26, 28]. The role of PEVs in promoting vascular endothelial cell proliferation, a key driver of angiogenesis, has been well established. Tang *et al.* showed that PEV enhanced the pro-angiogenesis of adipose-derived stem cells *in vivo* and *in vitro* and EVs from umbilical cord blood have been found to enhance angiogenesis in a mouse model [65, 66]. In our study, PEV-treated HUVECs had superior wound closure ratios compared to the control. However, cell migration was significantly reduced in both KM-only and PEV- KM treated groups, suggesting that KM may counteract the pro-angiogenic effects of PEVs, especially when taken together with our finding that PEV-KM together is the most effective inhibitor of tubular formation in endothelial cells. KM has been found to inhibit angiogenic activity in VEGF-stimulated human umbilical vein endothelial cells via the downregulation of PI3K/AKT, MEK and ERK pathways and targeting VEGFR-2 [67]. Interestingly, when HUVECs were treated with the PEV-KM combination, the overall effect was anti-angiogenic despite the inherent angiogenic potential of PEVs alone. The combination therapy may provide a way to modulate and fine-tune the balance between the promotion and reduction of angiogenic activity in the eye as needed. Collectively, our data show that the PEV-KM combination has the potential to mediate the angiogenic and inflammatory responses in human endothelial cells.

We also found that the combined PEV-KM formula was able to reduce corneal inflammation and angiogenesis *in vivo*, which was evidenced by the quantification of proinflammatory and proangiogenic cytokines in mouse eyes subjected to chemical cauterization-induced CoNV. The combined formulation also has the potential to attenuate corneal fibrosis, which could enhance overall corneal health and improve outcomes for corneal transplantation, as CoNV often impairs transplant success. Our findings, which show reduced levels of inflammatory and angiogenesis-associated proteins in KM-treated eyes (both KM-only and PEV-KM), are consistent with the literature surrounding the actions of KM in the cornea. Jia and others evaluated kaempferol for its ability to ameliorate the prognosis of fungal keratitis, a leading cause of corneal blindness via excessive inflammation in the cornea [68]. In their study, kaempferol was able to reduce immune cell infiltration and attenuate the expression of inflammatory factors such as TNF-α and IL-1β *in vitro* and *in vivo*. We observed that PEV may have a role in modulating fibrosis, typically during the corneal healing process. Specifically, α-SMA – a fibrotic marker – was significantly reduced in mouse eyes treated with PEVs, whether alone or in combination with KM, compared to those treated with PBS or KM alone. Our finding aligns with previous research showing that EVs can downregulate the onset of myofibroblasts (which express α-SMA and deposit collagen) through the inhibition of TFG-β2/SMAD2 pathways [69].

This study has certain limitations that warrant further investigation. Further work may include the use of cultured human corneal endothelial cells for *in vitro* evaluation as they may better reflect the *in vivo* cellular responses to PEV-KM compared to HUVECs. Further to this point, our *in vitro* experiments assessed the ability of PEV-only, KM-only and PEV-KM to penetrate a monolayer of HUVECs, while the real corneal environment is composed of many layers with different material properties. Additionally, PEVs are derived from donated material, meaning that there may be natural variability in the composition, stability and therapeutic molecular signatures of PEVs extracted from platelets. Finally, whilst we observed a reduction in inflammatory and angiogenic molecular signalling, we have not yet elucidated the exact mechanism by which this occurs. Future studies should aim to delineate the specific molecular pathways involved and assess the potential synergistic interactions among the tested compounds.

For the first time, our study demonstrates that the PEV-KM formulation effectively mitigates inflammatory responses and pathological angiogenesis in a mouse model of chemical cauterization- induced CoNV. The PEV-KM combination demonstrates a synergistic effect: PEV alleviates inflammation and supports wound healing, while KM reduces angiogenic stimulation. This synergy enhances the therapeutic efficacy of PEV-KM by utilizing PEVs to deliver KM directly to the target tissue, improving anti-angiogenic and anti-inflammatory outcomes. The therapeutic effects observed in this CoNV mouse model suggest the feasibility of PEV-KM as an effective agent for treating angiogenesis diseases in the eyes, especially corneal neovascularization, by topical therapy, such as eye drops. The translational applicability is reinforced by the fact that PEVs are a biological material that can be readily produced from clinical-grade platelet concentrates or autologous platelet donations [64].

## Acknowledgements

This work was supported by the integrated Research Grant in Health and Medical Sciences from the National Health Research Institute, Taiwan (NHRI-EX1109-10933SI, NHRI-EX110-10933SI, and NHRI-EX111-10933SI). GSL was supported by grants from the National Health and Medical Research Council of Australia (1185600 and 1123329). The Centre for Eye Research Australia receives Operational Infrastructure Support from the Victorian Government. The authors acknowledge the technical support provided by the TMU Core Facility- the Imaging Core for the TEM and confocal microscope. The authors also thank the TMU animal facility for assisting with the IVIS examination and the Taipei Blood Center for supplying the platelet concentrates to isolate the PEVs.

## Author contributions

Conceptualization- GS Liu, T Burnouf and CL Tseng. Methodology- GS Liu, T Burnouf and CL Tseng. Formal Analysis- GS Liu, HA Chen, CY Chang and CL Tseng. Investigation- HA Chen, CY Chang, YH Yang, YY Wu, YJ Chen, L Delila, and EH Hsieh. Resources- IC Lin, T Burnouf and CL Tseng. Writing (Original Draft)- GS Liu, HA Chen, T Burnouf and CL Tseng. Writing (Review & Editing)- GS Liu, HA Chen, A Widhibrata, T Burnouf and CL Tseng. Project Administration- CL Tseng. Funding Acquisition- GS Liu and CL Tseng. Project Supervision- GS Liu, T Burnouf and CL Tseng.

## Supplementary Data

**Figure S1.**
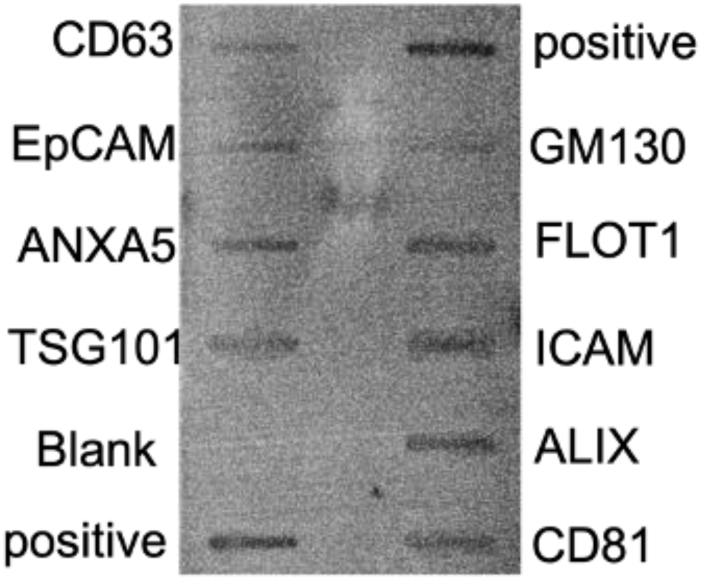
Images showing human-specific antibody arrays after incubation with PEVs. A total of 50 μg of protein was loaded onto the array. The bands observed from emitted light correspond to the presence of bound antibodies detecting EV markers such as CD63, ANXA5, TSG101, ALIX, CD81, FLOT1 as well as cell adhesion protein markers ICAM and EpCAM, and light expression of Golgi marker GM130.

**Figure S2.**
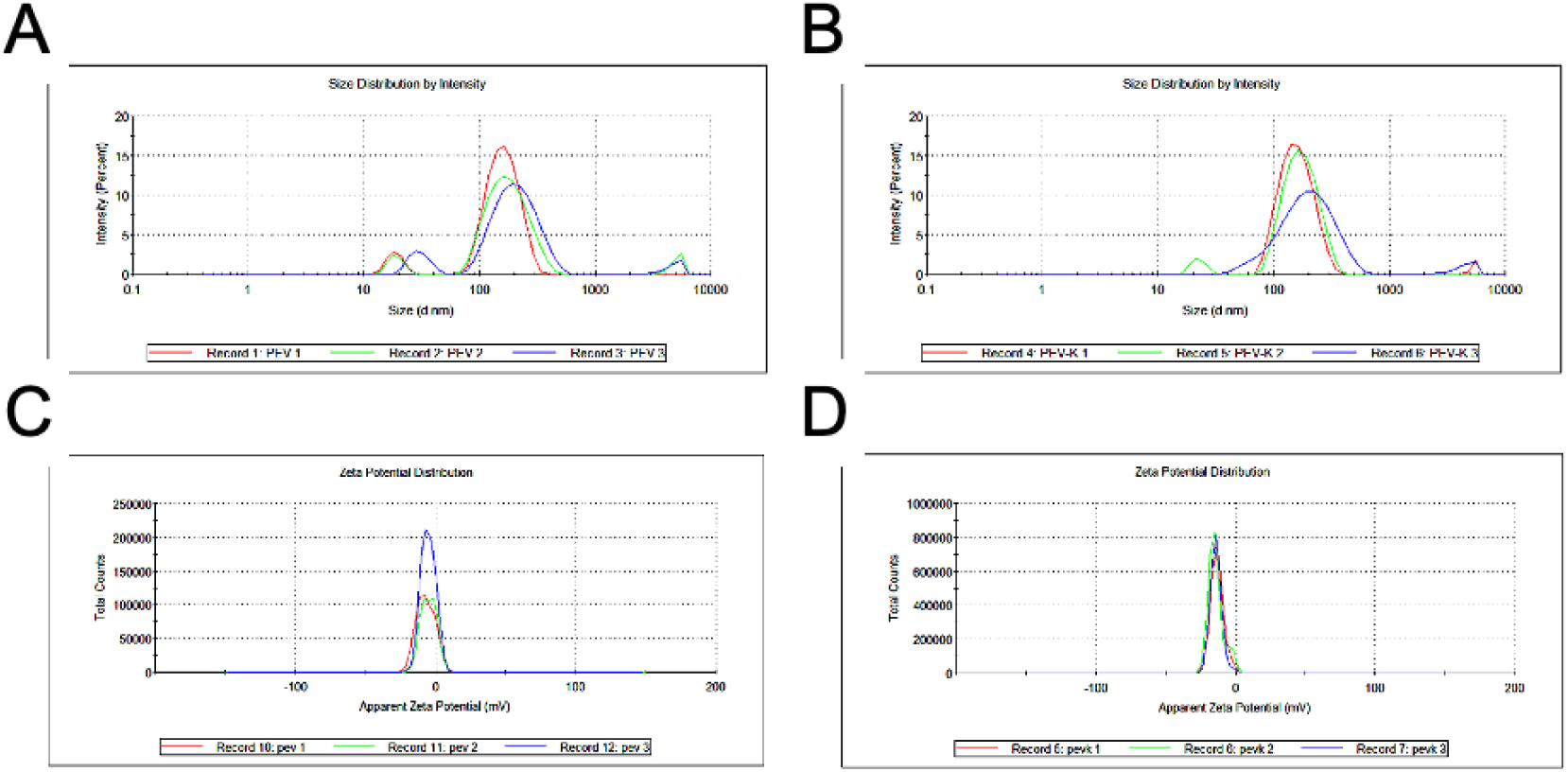
DLS result of PEV and PEV-KM. The particle size distribution of (A) PEV and (B) PEV- KM, as well as the peak zeta potential results of (C) PEV and (D) PEV-KM.

**Table S1.**
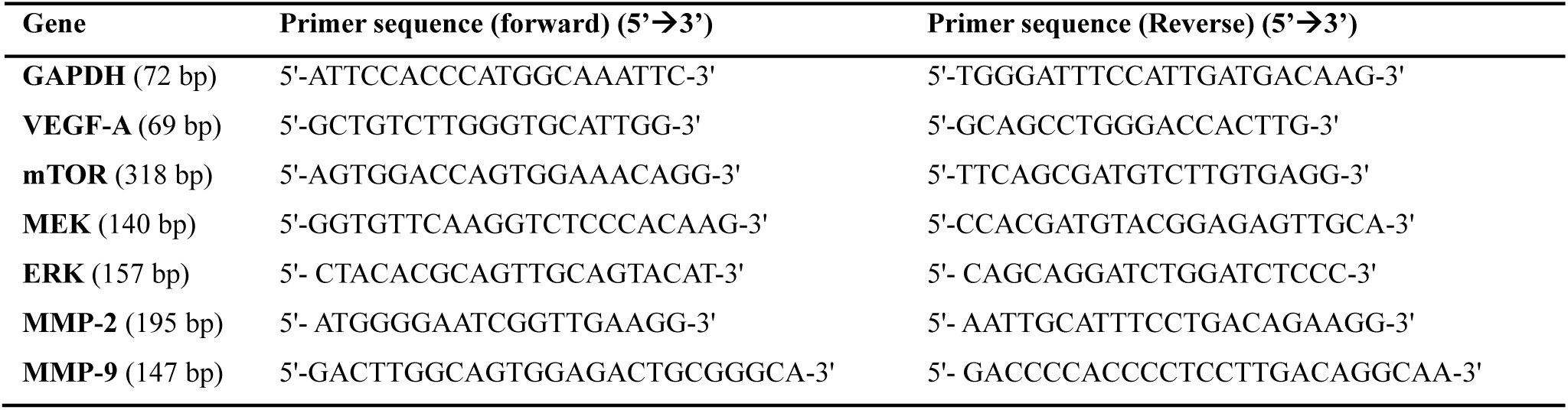
Primer sequence of angiogenetic markers for RT-qPCR.

**Table S2.**
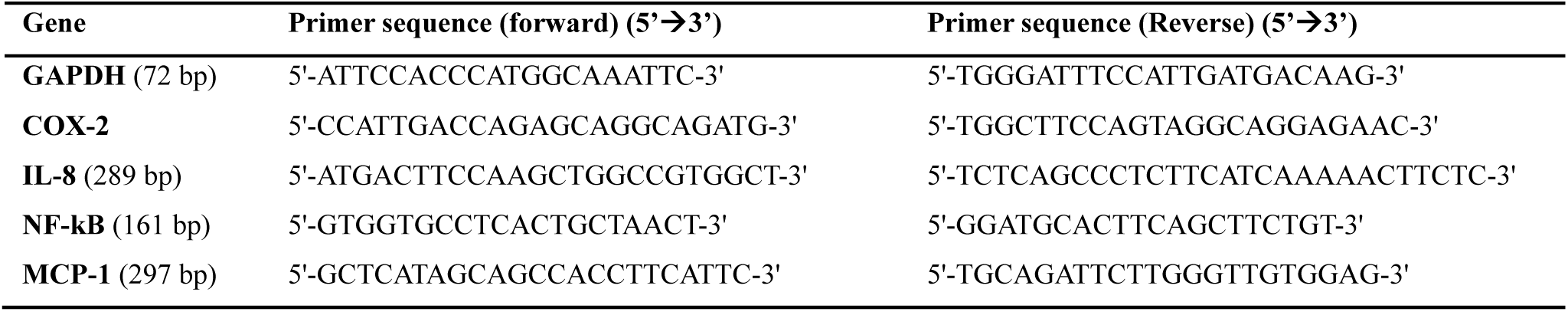
Primer sequence of Inflammatory markers for RT-qPCR.

